# AFM based elasticity of intestinal epithelium correlate with barrier function under drug action

**DOI:** 10.1101/761627

**Authors:** H. Tejeda-Mora, L. Stevens, M. Gröllers, A. Katan, E. van de Steeg, M. van der Heiden

## Abstract

Over the past few years, atomic force microscopy (AFM) has developed as a mature research tool for measuring the nanomechanical properties of tissue, cells and biological structures. The force spectroscopy mode of AFM allows the local elasticity of biological samples to be measured. The mechanical properties of cells are highly affected by homeostatic changes observed during disease. In the case of the intestine, the aetiology for some conditions is still unclear. To improve the clinical translation of pre-clinical models, a new and different approach could be to study cellular behaviour in health and disease from a mechanical point of view. Specifically, knowledge of changes in epithelial membranes in response to drugs is useful for interpreting both drug action and disease development. Here, we used human intestinal Caco-2 cells as a first step to record epithelial membrane elasticity measurements at the nanoscale using AFM. Three different drugs were selected to influence intestinal epithelium integrity by specifically targeting different functional aspects of the membrane, such as permeability and support. Results indicate a relationship between measured cell elasticity and cell viability markers, such as cellular toxicity and membrane barrier functions. Our work represents a proof-of-concept that cells suffer a particular change in elastic properties depending upon the mechanism of action of an applied drug. The following may provide an efficient approach for diagnosing intestinal pathologies and testing drugs for clinical use.

**STATEMENT OF SIGNIFICANCE:** We present evidence that epithelial membrane suffers a particular change in elastic properties depending upon the mechanism of action of an applied drug. These changes can be monitored over time using AFM technology and may provide an alternative and efficient approach for diagnosing intestinal pathologies and testing drugs for clinical use.

## INTRODUCTION

Atomic force microscopy (AFM) is a multifunctional technique used to investigate cellular responses with nanometre resolution under near-physiological conditions (1,2). The possibility of studying dynamic structural variations in the cell membrane is turning AFM into a relevant tool for developing models of the physical responses of cells to environmental stimuli (3-5). For this purpose, it is critical to be able to collect highly sensitive and selective mechanical data from the cell membrane in combination with techniques that provide information about the biology of the cell membrane. Recently, several studies have shown a relationship between several diseases and irregular cell mechanics. For instance, in cancer, at the tissue level, tumours were found to be relatively stiff (6,7), while at the cellular level, increased metastatic efficiency was found to be correlated with reduced stiffness (8,9).

The human intestinal epithelium is a monolayer of cells predominantly composed of enterocytes, but enteroendocrine cells, Goblet cells and M cells are also present. These cells form a monolayer sealed by epithelial junctions, which, among other functions, ensure barrier function and homeostasis (10,11). The intestinal tract is characterized by the formation of villi structures, and a remarkable characteristic of the enterocytes lining the villi is the presence of microvilli on the apical cell surface. Microvilli provide a large absorptive surface, which facilitates the absorption of nutrients, water and ions but also constitutes an effective barrier against undesirable substances (11-13). These characteristics make the intestinal epithelium an effective selective filter in organisms.

The functional behaviour of this selective filter, also referred to as barrier function, has been directly related to the viability of the organism (10,14). Examples of phenomena that can affect both are bacterial imbalance, defects in the epithelial barrier or/and immune regulation mechanisms (15-17). All these subsequently lead to disease and the development of inflammatory responses. Nevertheless, there is a gap in the understanding of the mechanisms involved in the transition to disease (18), thus hindering the treatment of several intestinal diseases. To better understand such diseases, an in-depth understanding of dynamic mechanical processes and functions is required. Here, we propose that monitoring mechanical properties at the nanoscale can be used to determine the viability of intestinal cells. In addition, fluctuations in the mechanical properties of the cells can be used to further understand the mechanism of action of membrane-targeting drugs.

In this study, human epithelial colorectal adenocarcinoma (Caco-2) cells, mimicking the intestinal epithelial membrane, were used to quantify changes in cell elasticity due to drug action. The Caco-2 cell line offers a good model for studying the intestinal epithelium because the phenotype and absorptive functions of Caco-2 cells approximate those of enterocytes (19,20). By applying three different membrane-targeting drugs, we recorded a varied response in cell elasticity using AFM technology. Our work demonstrates how intestinal epithelial cell membrane elasticity could serve as a bridge between the diagnosis and treatment of a disease. In the future, characterization of the cell membrane elasticity during drug action could be used to improve disease progression and drug development.

## MATERIALS AND METHODS

### Cell line and growth medium

Caco-2 cells (ATCC HTB 37) were purchased from the American Type Culture Collection (ATCC, Wesel, Germany). Caco-2 cells were routinely maintained in Dulbecco’s Modified Eagle Medium (DMEM, Gibco, Paisley, Scotland) with 10% foetal bovine serum (Gibco, Paisley, Scotland), 1% non-essential amino acids (Gibco, Paisley, Scotland), and 25 mg/ml gentamycin (Gibco, Paisley, Scotland). The cells were kept at 37 °C in a humidified atmosphere in the presence of 5% CO_2_. The medium was changed every 3 days.

### Membrane-targeting drugs

Three different conditions that affect cell integrity were tested. These conditions were studied in terms of mechanics and cell viability, and were triggered via the administration of drugs directly on the cell monolayer. During drug selection, drugs with clinical relevance were adopted. Two of these drugs, patulin and dextran sulphate sodium (DSS), are related to intestinal abnormalities. All selected drugs have an onset of action no longer than seven hours. The doses used are detailed in Table 1.

**Table 1.** Drugs used to trigger cell membrane damage; the clinically relevant concentration for each drug is marked in bold; concentrations used for AFM scanning are underlined.

| Drug | Doses | Reference |
| --- | --- | --- |
| Patulin | 1, 10, **50**, <u>100</u> μM | (28) |
| MCD | 2, **10**, 20 mM | (30) |
| DSS | 0.1, 1, **5** and <u>10</u> (w/v%) | (29) |

Patulin (Sigma-Aldrich Chemie B.V., Zwijndrecht, the Netherlands) is considered a toxic dietary compound that is secreted by several moulds. Patulin is generally present in spoiled fruits and has been reported to be toxic to bacteria, mammalian cell cultures, and other animals (20,21).

Cholesterol is a structural lipid present in all cells. Epithelial junction proteins depend on cholesterol-rich lipid domains (24). Here, we used methyl-β-cyclodextrin (MCD, Sigma-Aldrich Chemie B.V., Zwijndrecht, the Netherlands) to test the change in membrane elasticity during cholesterol uptake from the cell membrane. DSS (M_r_ 40,000, Sigma-Aldrich Chemie B.V., Zwijndrecht, the Netherlands) is a clinical chemical used to induce intestinal bowel disease (IBD). DSS is generally used to study the pathology of IDB. It has been reported that the cytotoxic potential of DSS depends on its molecular weight (29).

### Sample preparation

Caco-2 cells were cultured on polycarbonate 0.4-µm pore-size permeable filter supports (Corning Transwell Cambridge, MA, United States) for 21 days. Prior to AFM and cell viability measurements, the cell monolayer was gently rinsed with phosphate-buffered saline (PBS). After checking for a fully covered monolayer, the filter inserts were exposed to fresh, pre-warmed (37 °C) dose solutions containing the compound of interest.

The drug compound at the desired concentration and a membrane integrity marker for permeability reference measurements (FD4, 50 µM) were added to the apical side of the insert. Only the dose solutions containing MCD were added to both the apical and basolateral compartments of the insert. For AFM measurements, only one dose was tested (Table 1).

Samples used for AFM measurements and cell viability measurements came from the same cell batch but were measured independently. For all AFM measurements, the polycarbonate membrane with attached Caco-2 cells was carefully cut around the edges with a scalpel and clamped in preparation for AFM imaging (Fig. 1). Immediately after mounting the sample, the cells were exposed to medium containing the drug of interest until the space between the sample and MTFML-V2 probe holder (Bruker, Santa Barbara, CA) was filled.

**Figure 1.**
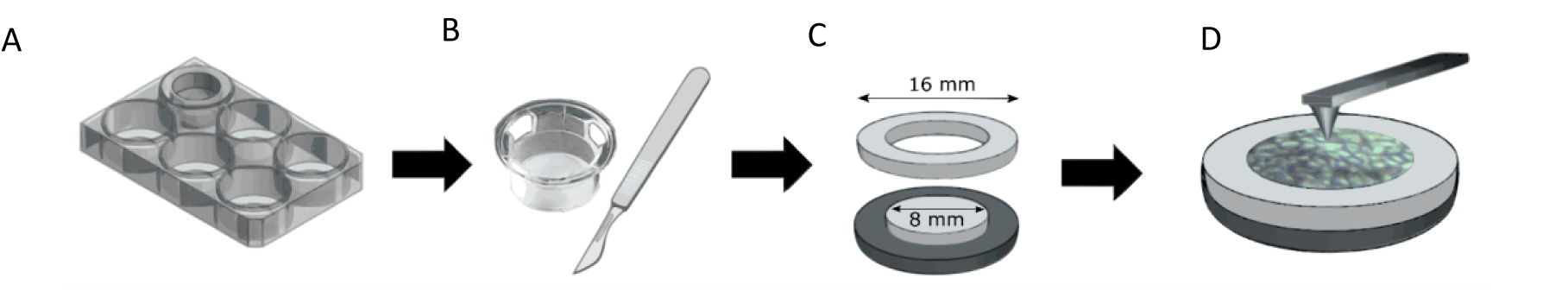
Schematic of the procedure for AFM sample preparation. A) Permeable membrane inserts used for Caco-2 cells. B) For each insert, the membrane was cut. C) Setup used for AFM imaging. The base was made of iron for magnetic attachment to the AFM sample holder. The upper part of the base and ring were made of nylon. The cut membrane was clamped between the two pieces. D) The sample was placed for AFM analysis in liquid environment.

### AFM setup

The AFM measurements were performed using an AFM MultiMode 8 system (Bruker, Santa Barbara, CA) with silicon nitride cantilevers. C-MLCT probes (Bruker, Santa Barbara, CA) with a nominal spring constant of 0.01 N/m and a nominal tip radius of 20 nm were used. High lateral resolution from the sharp probes allowed to obtain precise point measurements from the cell membrane. Deflection sensitivity and cantilever stiffness were calibrated before the measurements. Force-volume data was recorded in liquid at room temperature. Recorded data per sample covered an area of 20×20 µm^2^ (roughly the area of 4-8 cells). A ramp size of 5 µm was used. The ramp rate was kept between 1-1.5 Hz. The trigger was set relative to a threshold of 1.5 nN. This allowed to record indentations close to 1 µm and still be in the linear regime for elastic materials, allowing a Hertz contact model for extraction of the elastic parameters (31).

### Force-volume analysis

The force-volume files were processed using NanoScope Analysis software (Bruker, Santa Barbara, CA) to obtain cell membrane elasticity values. Young’s modulus was calculated using 60-85% of the indentation section of the force-distance curve (Fig. 2A). The contact point from each curve was checked by visual inspection. Force-distance curves were excluded from the analysis when the contact point could not be determined due to false force-distance curve records.

**Figure 2.**
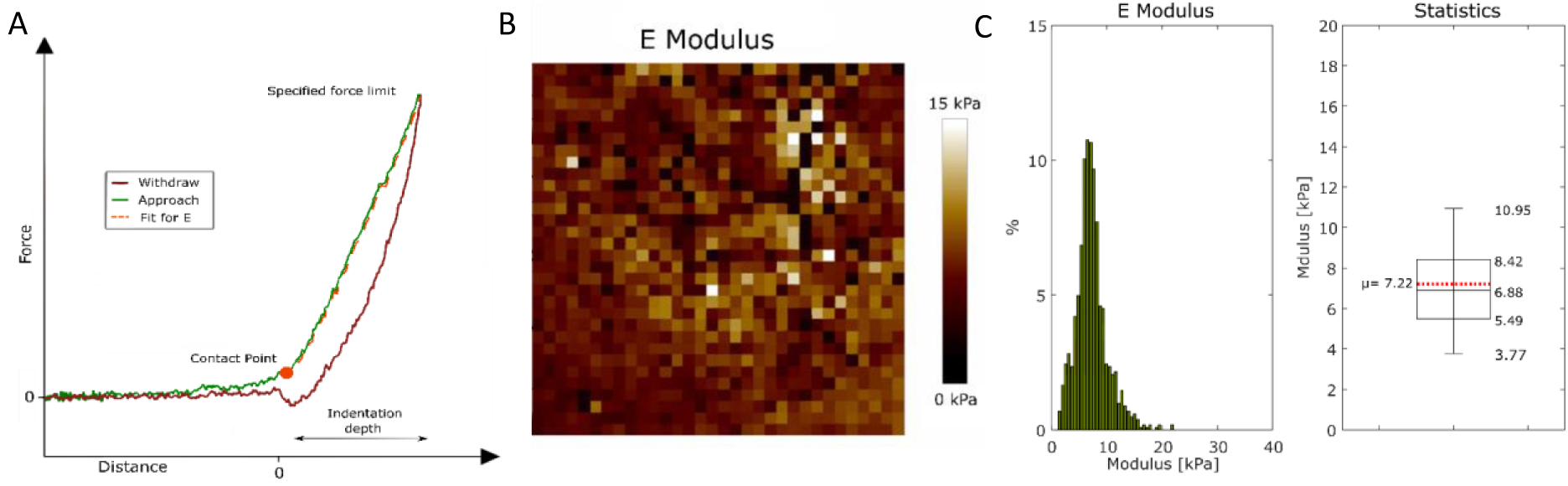
Caco-2 cell membrane elasticity. A) Characteristic force-distance curve recorded as point measurements in force-volume mode. B) Map of Caco-2 monolayer modulus over an area of 20×20 µm^2^. C) Statistical analysis. Left, histogram of the distribution of the membrane E modulus. Right, box plot with interquartile range. The mean is shown in red. Whiskers show 10-90% of the data.

### Cell barrier integrity via trans-epithelial electrical resistance (TEER) measurements

The viability of the Caco-2 cell monolayers was assessed by measuring three different viability markers after drug application. The TEER of the monolayers was measured using a Millicell ERS-2 epithelial volt-ohmmeter (Millipore, Billerica, MA). The integrity of the Caco-2 barrier was considered unaffected when the TEER values remained stable more than 100 Ω/cm^2^ above the background. Monolayers with values less than 100 Ω/cm^2^ above the background were excluded from the measurements. TEER measurements were also used as a membrane integrity marker after cells were exposed to the drugs.

### Barrier permeability determination using FD4

The integrity of the Caco-2 cell monolayer barrier was assessed by measuring leakage of the paracellular transport marker fluorescein isothiocyanate–dextran MW 4000 (FD4, 50 µM) (Sigma-Aldrich Chemie B.V., Zwijndrecht, the Netherlands) over time into the basolateral compartment. The intestinal barrier was considered intact when the FD4 leakage was less than 1.0%. After sampling, the original volume from the apical compartment was refilled with pre-warmed dose solution. The samples were analysed using a BioTek Synergy multimode microplate reader (BioTek Instruments, Winooski, VT, United States) at an excitation wavelength of 485 nm and an emission wavelength of 528 nm.

### Lactate dehydrogenase (LDH) activity cytotoxicity assay

Cell death was evaluated by measuring the activity of cytoplasmic enzymes released by damaged cells. Here, LDH was used. LDH activity was determined using an LDH kit (Sigma-Aldrich/Roche, Zwijndrecht, the Netherlands) following the supplier instructions. Briefly, samples were collected from the apical compartment (50 µL) at every time point. After sampling, the original volume from the apical compartment was refilled with pre-warmed dose solution. All obtained samples were incubated protected from light with the reaction mixture (50 µL) from the LDH kit for 30 min. Then, 1 M HCl (20 µL) was added to stop the reaction. The absorbance at 490 nm was recorded using a BioTek Synergy multimode microplate reader.

### Statistical analysis

A minimum number of three biological replicates were analysed to ensure reproducibility for all experiments. For each experimental group, the mean and standard deviation were calculated. Manual validation of the AFM data resulted in the exclusion of less than 5% of the data. The main cause was attributed to artefacts during measurements, which caused false force-distance curve recordings. Box plots show the median, interquartile range (25-75%) and 10-90% of the data (whiskers), including the mean (marked in red). The mean and standard deviation were calculated separately. Data were analysed by one-way ANOVA. P-values less than 0.01 were considered statistically significant. Comparisons were made between groups to validate significant differences between mean values.

## RESULTS

To verify the validity of AFM-based cellular elasticity measurements, monolayers of Caco-2 epithelial cells were used. AFM was performed on three cell monolayers (Fig. 2B, C). The elasticity of the cell membrane was recorded as 7.2 ± 4.7 kPa (mean ± std. dev.). Individual cells were not distinguishable on the modulus map (Fig. 2B) due to the dense microvilli on the cell membrane. The TEER of healthy Caco-2 monolayers was shown to be higher than 100 Ω/cm^2^ above the background, confirming cell confluency.

### Cell membrane elasticity changes over time due to membrane damage

To investigate the relationship between disease-mimicking conditions and the mechanical properties of the cell membrane, three different drugs (patulin, MCD, DSS) were applied to the cell monolayers. Cell viability and barrier integrity were monitored for six hours. Similarly, cell monolayers were probed using AFM. To exclude the influence of sample-tip interactions on cell membrane elasticity, untreated cells were probed every hour for six hours. No significant changes were observed between the first and the following measurements (all p-values > 0.18), indicating that the exposure of cells to ambient environments and constant probing by AFM do not affect cell membrane elasticity (Fig. 3). The average value of all reference measurements is indicated by the blue dashed line in the box plots from Figures 3-7. Caco-2 monolayers were monitored for six hours after drug exposure to evaluate cell integrity and membrane elasticity in response to each drug independently. We found that control TEER and FD4 leakage measurements showed a slight decrease over time. Such decreases can be explained by the lack of optimal O_2_ and CO_2_ conditions during sampling. However, LDH (cytotoxicity marker) release was stable for untreated cells, indicating an intact cell membrane. In addition, all applied drugs caused a significant change in cell membrane elasticity after four hours (Fig. 4-6). The change in membrane elasticity was compared to biological changes in membrane functions to evaluate the effect of the applied drug from a mechanical perspective.

**Figure 3.**
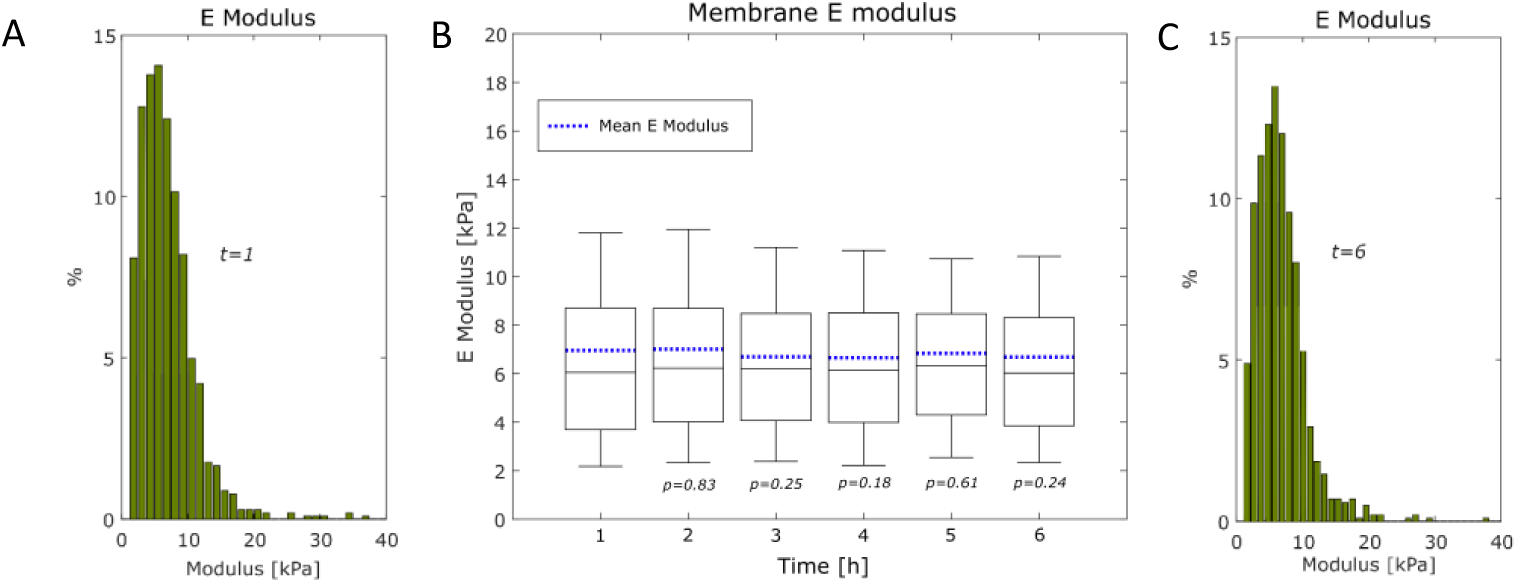
Elasticity of Caco-2 cell membrane over six hours. A, C) Statistical analysis. Histograms of the distribution of the membrane E modulus at t=1 and t=6 hours for comparison. B) Membrane elasticity. The mean is marked in blue for each group.

**Figure 4.**
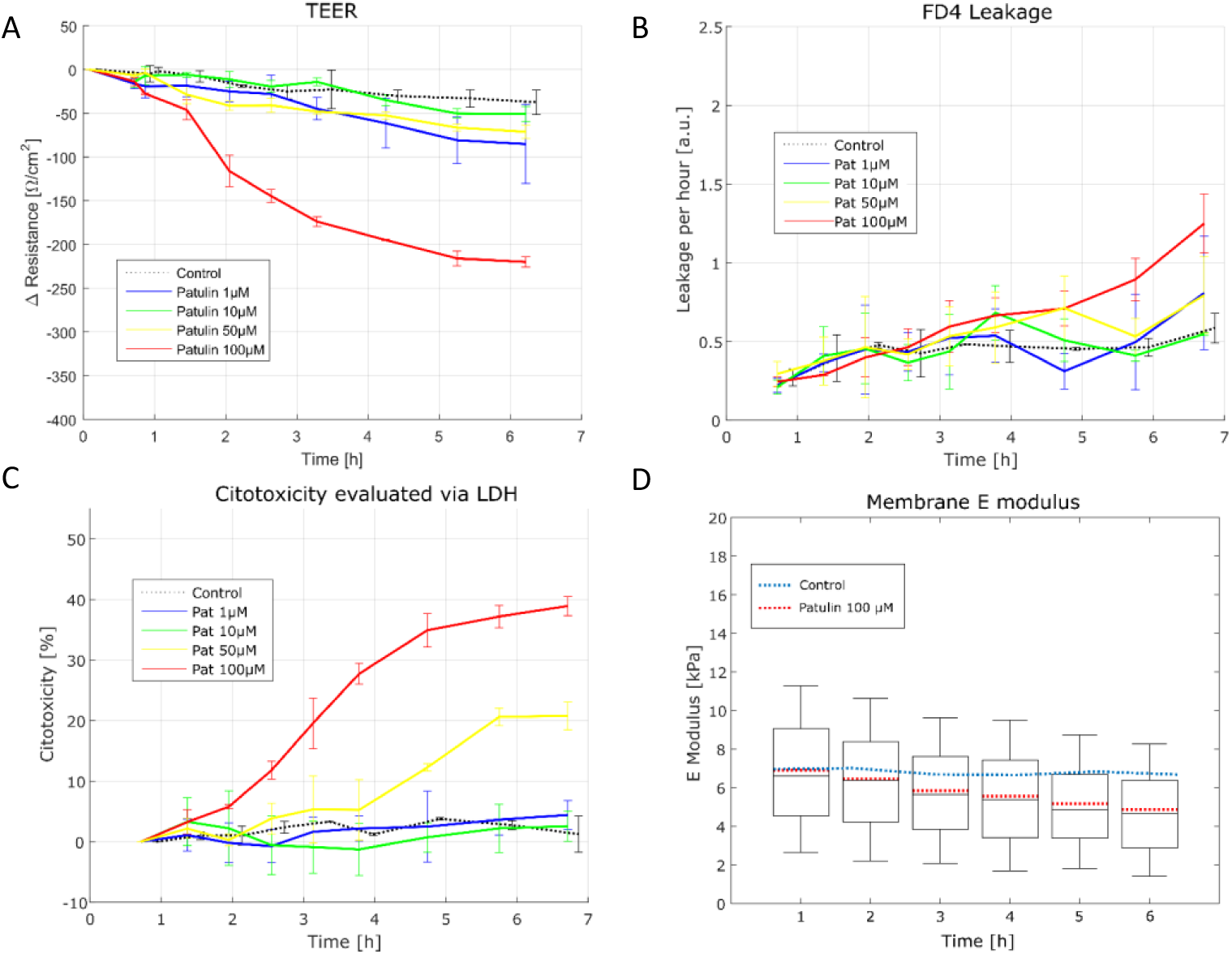
Effects of patulin on Caco-2 cells. A) TEER measurements expressed as the relative change in resistance for each concentration. The red box marks the time point and concentration used for AFM measurements. B) Permeability of the cell membrane measured by FD4. C) Cell membrane damage expressed as a percentage compared to lysed cells. D) Effect of patulin on membrane elasticity over six hours (100 mM). The mean is marked in red for each group. The blue line shows the mean values of untreated cells.

**Figure 5.**
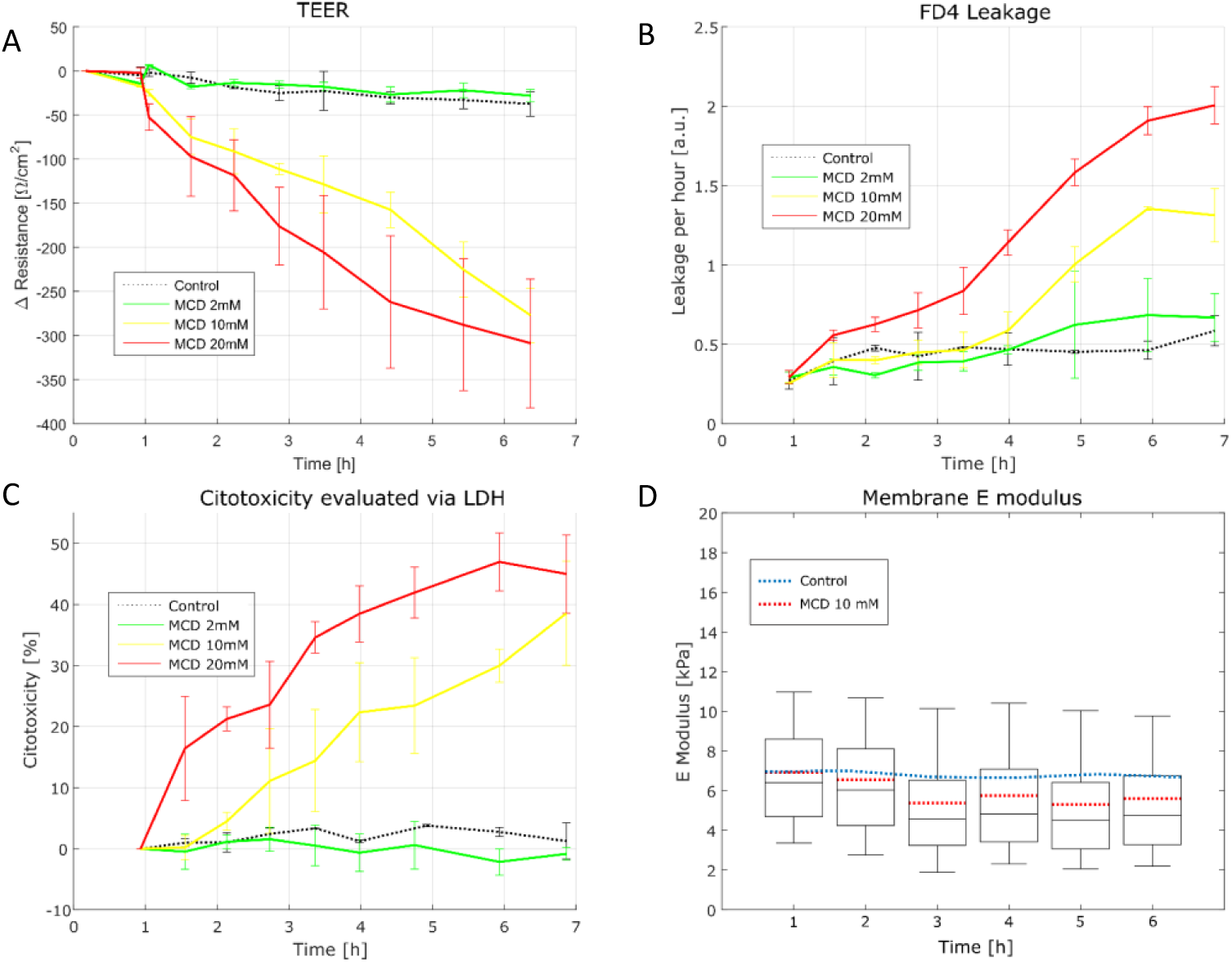
Effects of MCD on Caco-2 cells. A) Measurements are expressed as the change in resistance for each concentration. The red box marks the time point and concentration used for AFM measurements. B) Permeability of the cell membrane measured by FD4. C) Cell membrane damage is expressed as a percentage compared to lysed cells. D) Effects of MCD on membrane elasticity over six hours (10 mM). Statistical parameters are shown as box plots for each hour. The mean is marked in red for each group. The blue line shows the mean values of untreated cells.

**Figure 6.**
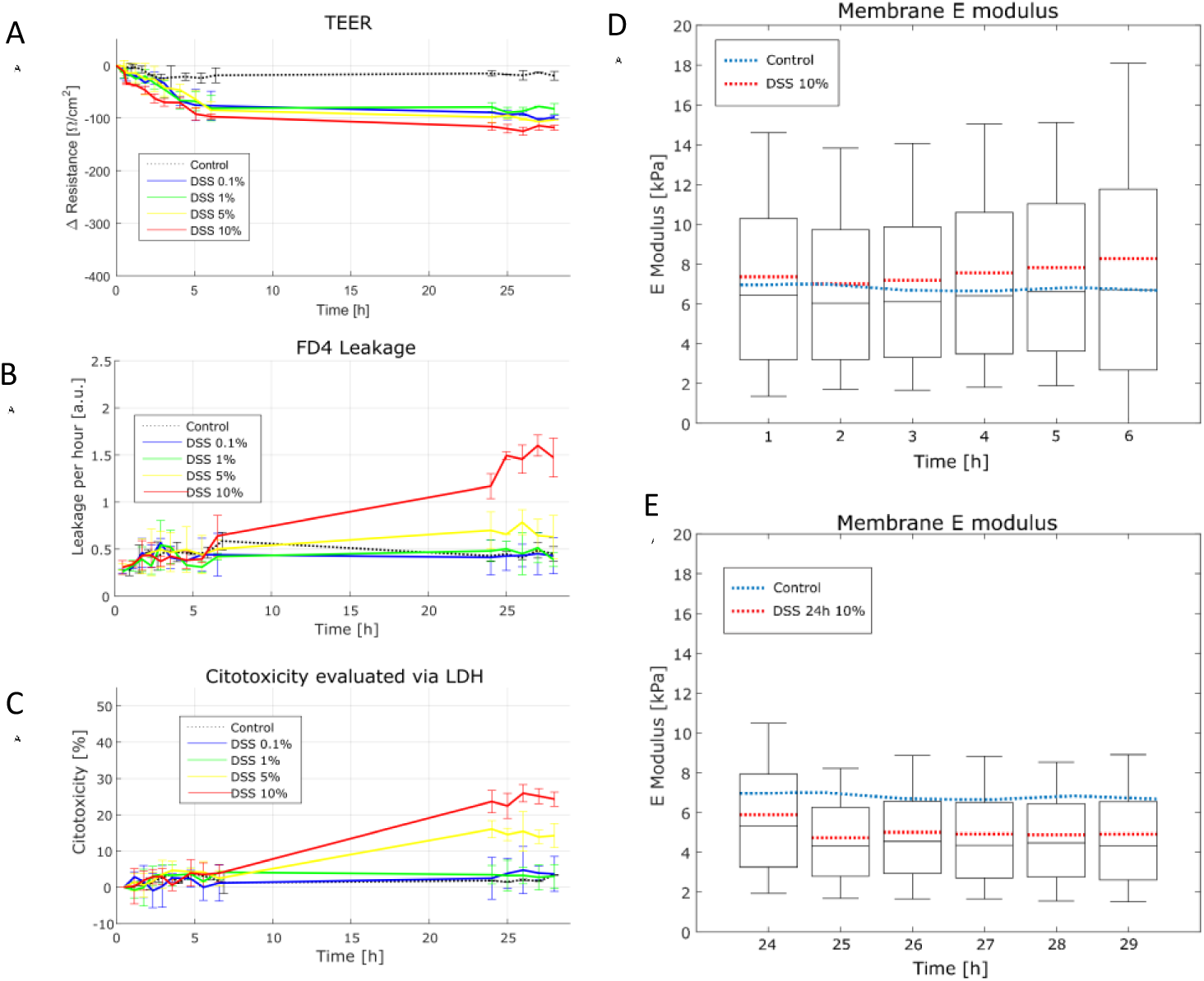
Effects of DSS on Caco-2 cells. A) TEER measurements are expressed as the change in resistance for each concentration. The red box marks the time point and concentration used for AFM measurements. B) Permeability of the cell membrane measured by FD4. C) Cell membrane damage is expressed as a percentage compared to lysed cells. D-E) Effect of DSS on membrane elasticity over six hours (10% w/v). Statistical parameters are shown as box plots for each hour. The mean is marked in red for each group. The blue line shows the mean values of untreated cells.

### Patulin causes a reduction in the membrane elastic modulus

Patulin is a mycotoxin that affects the cell membrane by interfering with epithelial junctions due to its strong affinity for sulfhydryl (SH) groups present in membrane proteins and in the cell membrane itself (22,23), and it was tested at different concentrations for 6 hours. Figure 4A shows a significant decrease over time in the electrical resistance of the membrane of cells treated with patulin at 100 µM. All other concentrations showed a nosignificant decrease over time. Patulin caused no significant disruptions in the tight junctions, as reflected by low FD4 leakage across cellular monolayers for all concentrations tested (Fig. 4B). Nevertheless, patulin damaged the cell membrane at the higher concentrations of 100 and 50 µM, as shown by the release of LDH from the cells (Fig. 4C). Our cell viability results are in accordance with those of McLaughlin et al. showing that patulin causes apoptosis and a decrease in intestinal barrier function (21-23, 24).

In a parallel experiment, Caco-2 cells treated with the highest concentration of patulin (100 µM) were probed using AFM for six hours. The membrane E modulus was reduced from 7.2 ± 4.7 kPa to 4.9 ± 3.6 kPa after six hours (Fig. 4D) compared to untreated cells (p<0.001). This decrease is expected to be due to membrane damage and is in good agreement with the TEER, FD4 leakage and cytotoxicity measurements.

### Cholesterol depletion by MCD reduces cell membrane elasticity

Cholesterol depletion causes an imbalance in the function of junction proteins and enhances epithelial permeability. MCD has been reported to cause a rapid reduction in cholesterol by extraction through direct contact with the plasma membrane. MCD does not bind to the plasma membrane or enter the cell, resulting in no interference with metabolic activity (25,26). Figure 5 shows the effects of 2, 10 and 20 mM MCD after 6 hours; the 2 highest concentrations of MCD (10 and 20 mM) caused a significant change in all reference measurements over time (Fig. 5 A-C). The higher the amount of cholesterol taken from the cell membrane, the more the cell barrier is damaged, as indicated by all integrity parameters (Fig. 5A-C). Significant damage to the cell membrane and protein junctions was also observed (Fig. 5 B, C), confirming that at higher concentrations, MCD is highly toxic to Caco-2 monolayers.

Because cholesterol is an important building block of the cell membrane and provides support to tight junctions, its depletion by MCD was expected to cause a decrease in cell membrane elasticity. This was tested at a concentration of 10 mM using AFM. The obtained membrane elasticity is displayed in Figure 5D. A significant decrease in the elastic modulus (5.8 ± 3.2 kPa) of the membrane was observed compared to reference cells (p<0.001), justifying the hypothesis that cholesterol aids in maintaining cell membrane rigidity.

### DSS decreases the membrane elastic modulus

DSS is applied as a chemical inducer to mimic IBD in animal models. The interaction of sulphate groups with the cell membrane has been reported to produce tissue inflammation and an increase in membrane permeability (26). The damage caused by DSS is believed to occur in epithelial junctions, resulting in the dissemination of inflammatory intestinal contents into underlying tissue. We applied DSS at concentrations of 0.1, 1, 5 and 10% (w/v) for an exposure time of six hours. Figure 6A shows the TEER values of Caco-2 monolayers exposed to DSS, demonstrating a small decrease in the monolayer resistance over time for all DSS concentrations. Unexpectedly, the leakage of FD4 across the cell membrane did not change over the six hours after treatment with DSS (Fig. 6B). Additionally, DSS did not cause significant cell membrane damage during the first 6 hours after application, as illustrated in Figure 6C. However, after 24 hours, the cell membrane permeability and cytotoxicity markers showed significant changes for the two higher concentrations of DSS, indicating a delayed effect of DSS on the cell membrane.

Simultaneously, AFM experiments were performed to study the effect of DSS at 10% (w/v) on membrane elasticity. DSS on its own caused variation in the membrane elasticity after the first hour (Fig. 6D). The force-volume data revealed a significant increase in the membrane elasticity compared to reference measurements. The average membrane E modulus (8.2 ± 4.3 kPa) showed a significant increase compared to healthy cells (Fig. 6D) (p<0.001). Here, the higher E modulus could have resulted from a cellular stress response to DSS. To validate the influence of DSS, additional measurements were performed 24-29 hours after exposure (Fig. 6E). Surprisingly, after 24 hours, DSS caused a significant decrease in membrane elasticity (p<0.001), which remained constant over the probed time (4.9 ± 2.9 kPa after 26 hours of application).

## DISCUSSION

Our work demonstrates how the nanomechanical properties of the cell membrane adapt to the environment and could be used as an additional non-invasive in vitro method for determining the mechanical and elastic adaptations of the cell membrane as a parameter of cellular stress. The relationship between the effects of drugs and the intestinal membrane elasticity was examined using Caco-2 monolayers as an intestinal model. The use of three clinically and experimentally relevant drugs furthered the understanding of how the mechanism of action of a drug affects the mechanical properties of the cell membrane. We also demonstrated the possibility of monitoring the cell membrane over time under near-physiological conditions without influencing cell viability due to AFM sampling. Thus, AFM can potentially be used as a non-invasive tool for studying the effects of compounds (e.g., drugs) on cells and cell membrane behaviour.

During AFM probing, it is important to closely mimic cell culture conditions to reduce possible sources of noise that could influence measurements. In this study, we attempted to do this by performing in vitro experiments in a liquid environment and under static controlled conditions. In subsequent experiments, the use of a flow cell could improve not only the nutrient and drug supply but also waste removal, allowing longer probing times.

The force-distance curves obtained from all samples presented the characteristic behaviour of viscoelastic materials, as indicated by the nonlinearity of the indentation section of the curve (Fig. 2A). In this study, we specifically focused on extracting the elasticity of the cell membrane, and we did not calculate energy dissipation, relaxation, or the influence of the microvilli or cell membrane adhesion because these calculations require more complex analytical models. Therefore, we cannot absolutely confirm that drug action disrupts only cell membrane elasticity.

We decided to work using the force-volume mode with slow scanning rates (1-1.5 Hz) to allow the recording of force-distance curves from the membrane accurately in an automated way. Faster AFM modes exist, such as multifrequency modes, but cell membrane heterogeneity and the liquid environment challenge data collection in these modes. In addition, these modes show less adequate elasticity estimations due to the complexity of analysing harmonic data (31,32), making them impractical for potential clinical use. Another point to consider is the viscoelastic behaviour of cells. The recorded force-distance curves reveal information related to the membrane and cytoskeletal components that are deformed under stress. At higher frequencies, membrane components show rigid body behaviour (33). It would be interesting to observe the membrane behaviour at different frequencies, avoiding very high frequencies that may be irrelevant for physiological timescales (> 100 Hz).

Although genetic and metabolic changes are correlated with many intestinal diseases, the aetiology of some diseases is not yet well understood, making diagnosis and treatment challenging; such diseases include IBD, Irritable bowel syndrome (IBS) and even cystic fibrosis (34-36). Previous work performed with different cellular models and other mechanobiological techniques has suggested different mechanisms linking intestinal diseases with diverse metabolic pathways (37,38). We propose that membrane elasticity can be used as an additional signature for measuring and monitoring cellular stress under various conditions. Here, we show that patulin and MCD cause a reduction in membrane elasticity over time, which could be explained by biological responses involving toxicity (LDH) and membrane barrier function (FD4 and TEER). In addition, we observed that cholesterol is crucial for maintaining cellular membrane viability but that the elasticity of the membrane does not decrease as much with MCD as with patulin or DSS. This finding supports the idea that cholesterol is essential for keeping epithelial junctions sealed but that the membrane elasticity is more susceptible to stress caused by a drug than to cholesterol depletion.

An unexpected response was observed when Caco-2 cells were exposed to DSS. During the first six hours, cell viability remained intact as determined by biological indicators (LDH, FD4 and TEER), but the cell membrane elasticity was significantly increased. Remarkably, this increased membrane elasticity was reversed after 24 hours of exposure, when a reduction in cell membrane elasticity was observed. This reduction in elasticity reflected a decrease in cell viability. We can only speculate about the reason for this change, but we think it was due to osmotic effects induced by DSS during the first hours of cellular interaction, resulting in an increase in membrane stiffness but no cell damage. Later, DSS permeated the cell monolayer, causing cytotoxic effects to the epithelium and a reduction in membrane elasticity. This delayed response may be linked to immunological or fibrolytic mechanisms, as suggested in (25).

Membrane damage can be measured in terms of its nanomechanical properties, improving prospects for disease diagnosis and drug development from a mechanical point of view. Besides, our results have provided the first milestone to achieve the monitoring functional behaviour of living cells, by probing the cell membrane. Future studies with patient biopsiesmaterial as primary cell cultures or tissues will probably identify additional aspects for specific conditions but are unlikely to change the overall conclusion that cell stress shapes the mechanical properties of the cell membrane.

## AUTHOR CONTRIBUTIONS

E.S., M.H., and H.T. designed the experiments. H.T., M.G., and L.S. performed the experiments. E.S., A.K. and H.T. developed experimental protocols for sample preparation. M.H. and H.T. analysed related data. M.H. and E.S. supervised the research. H.T. wrote the manuscript.

## ACKNOWLEDGEMENTS

This research project was financially supported by the early research programmes for organs on a chip and 3D nanomanufacturing of the Dutch Research Institute for Applied Research, TNO.

